# An artificial host system enables the obligate parasitic plant *Cuscuta campestris* to grow and complete its life cycle *in vitro*

**DOI:** 10.1101/2021.06.21.449293

**Authors:** Vivian Bernal-Galeano, James H. Westwood

## Abstract

*Cuscuta campestris* is an obligate parasitic plant that requires a host to complete its lifecycle. Parasite-host connections occur via an haustorium, a unique organ that acts as a bridge for the uptake of water, nutrients and macromolecules. Research on *Cuscuta* is often complicated by host influences, but comparable systems for growing the parasite in the absence of a host do not exist. We developed an axenic method to grow *C. campestris* on an Artificial Host System (AHS). We evaluated the effects of nutrients and phytohormones on parasite haustoria development and growth. Haustorium morphology and gene expression were characterized. The AHS consists of an inert, fibrous stick that mimics a host stem, wicking water and nutrients to the parasite. It enables *C. campestris* to exhibit a parasitic habit and develop through all stages of its lifecycle, including production of new shoots and viable seeds. Phytohormones NAA and BA affect haustoria morphology, and increase parasite fresh weight and biomass. Gene expression in AHS haustoria reflect process similar to those in haustoria on actual host plants. The AHS is a methodological improvement for studying *Cuscuta* biology by avoiding specific host effects on parasite and giving researchers full control of the parasite environment.

## Introduction

Dodders are parasitic plants of the genus *Cuscuta* (Convolvulaceae) that can invade a wide range of dicot plants. *Cuscuta* spp. are characterized by their vining growth habit and lack of roots and expanded leaves. They use an organ unique to parasitic plants, the haustorium, to invade host stems to gain access to essential nutrients and water resources. *Cuscuta* spp. exhibit compromised or no photosynthetic activity (Van Der Kooij *et al.*, 2000), so they rely at various levels on their hosts as their only carbon source. Thus, their survival depends on parasitic features like the ability to find a host, produce functional haustoria and efficiently take up resources.

*Cuscuta* spp. are obligate parasites, so must have a host to complete their life cycles (Westwood *et al.*, 2010). The *Cuscuta* spp. life cycle comprises the stages seed, seedling, host colonization, haustoria development, vegetative growth, flower production, and fruit and seed set. *Cuscuta* seeds germinate on the soil without any chemical stimulant (Benvenuti *et al.*, 2005), and the thread-like seedlings must find a suitable plant host to avoid death due to exhausting its limited resources (Shimizu & Aoki, 2019). After the parasite has found a host, its shoot coils around the host stem, and the haustorium starts to develop from within the internal face of the coil. Coiling and haustorium formation respond positively to the exposure to enriched far-red light (700-800 nm) and tactile stimuli (Lane & Kasperbauer, 1965; Tada *et al.*, 1996). These stimuli provoke an increase in internal levels of the phytohormone cytokinin, leading to haustoria induction (Haidar *et al.*, 1998; Furuhashi *et al.*, 2011).

Haustorium development involves tissue dedifferentiation and redifferentiation, and can be divided into three phases: adhesive, intrusive, and conductive (Heide-Jørgensen, 2008). In the adhesive phase, epidermal cells redifferentiate into trichome-like cells that form a holdfast or adhesive disc (Vaughn, 2002). Holdfast cells secrete pectin and other compounds that mediate the adhesion of parasite tissues to host tissues. Meanwhile, meristematic cells in the inner cortex behind the holdfast generate the prehaustorium (Shimizu & Aoki, 2019).

During the intrusive phase, the prehaustorium becomes the haustorium (Heide-Jørgensen, 2008). The haustorium, formed by intrusive cells, elongates through the center of the holdfast and penetrates the host stem epidermis and cortex (Vaughn, 2003). Then terminal digitate cells, or searching hyphae, elongate to encounter host vascular elements. Initial haustoria formation is independent of host factors, however, they may be required for further differentiation of xylem vessels (Kaga *et al.*, 2020). Searching hyphae cells adjacent to host xylem elements become xylem hyphae, while those adjacent to sieve tubes become absorbing hyphae. Subsequently, cortex cells in the haustorium also differentiate into phloem or xylem elements as appropriate (Vaughn, 2006; Shimizu & Aoki, 2018; Shimizu *et al.*, 2018).

The conductive phase starts after searching hyphae differentiate to vascular conductive elements in the mature haustoria, completing the vascular connection between the host and the parasite (Heide-Jørgensen, 2008). The parasite becomes a strong sink competing with the host for resources as exemplified by *C. reflexa*, which obtains 99% of its carbon from the host (Jeschke *et al.*, 1994). Once the connections are established, the *Cuscuta* shoot resumes growth to produce new shoots that reach other host stems, making additional points of connection and ultimately produces flowers, fruits, and seeds.

Recent studies have described interesting features of the host-*Cuscuta* system including the reciprocal exchange of large amounts of macromolecules such as proteins (Liu *et al.*, 2020), mRNAs (Kim *et al.*, 2014), and microRNAs (Shahid *et al.*, 2018). Some of the exchanged microRNAs have been shown to function by affecting gene expression in the host plant (Shahid *et al.*, 2018). There is an urgent need to develop new methods to study *Cuscuta* in ways that can distinguish parasite processes from host processes in order to define the contributions of each to the interaction.

One approach to studying *Cuscuta* in the absence of a host is through tissue culture. Published methodologies exist for obtaining callus and shoot regeneration for *C. trifolii* (Bakos A. *et al.*, 1995; Borsics *et al.*, 2002), *C. europea* (Švubová & Blehová, 2013), and *C. reflexa* (Srivastava & Dwivedi, 2001), and floral induction for *C. reflexa* (Baldev, 1962). Other studies reported the production of haustoria by applying a mechanical pressure between two surfaces, along with a far-red light stimulus on shoots of *C. japonica* (Tada *et al.*, 1996; Furuhashi *et al.*, 1997), *C. reflexa*, and *C. campestris* shoots (Kaga *et al.*, 2020; Lachner *et al.*, 2020). These systems have certain advantages, but tissue culture does not allow investigations related to haustorial form or function, and none of them accurately reflect normal *Cuscuta* parasitic growth habit with production of new shoots and viable seeds.

Here we report the development of an axenic system to grow *C. campestris* on an artificial host that provides water, nutrients, and phytohormones to the parasite. We demonstrate that *C. campestris* growing in this Artificial Host System (AHS) exhibits all the stages observed during its typical lifecycle, including flower, fruit, and production of viable seeds. Functional characterization, including transcriptomic analysis, of the haustorial region of *C. campestris* growing in the AHS shows growth that is similar to that on a living host.

## Materials and methods

### Preparation of the artificial host system

The AHS was assembled using a magenta GA-7 plant culture box (Sigma) and a plastic autoclavable lamina (Polypropylene No. 5, 0.4 mm thick) cut to fit the bottom of the box and serve as a support to hold the artificial hosts (Fig.**1**, **S1**). The paper spindle from a cotton swab was used as the “host stem”. Each magenta box held four spindles. When *Cuscuta* seedlings were used as inoculum, 0.2ml microcentrifuge tubes were attached to the base of each spindle to hold the seedling. All the system components, magenta boxes, sheets with artificial hosts and aluminum foil strips (used to attach the plant to the artificial host) were autoclaved before assembly.

**Fig. 1.**
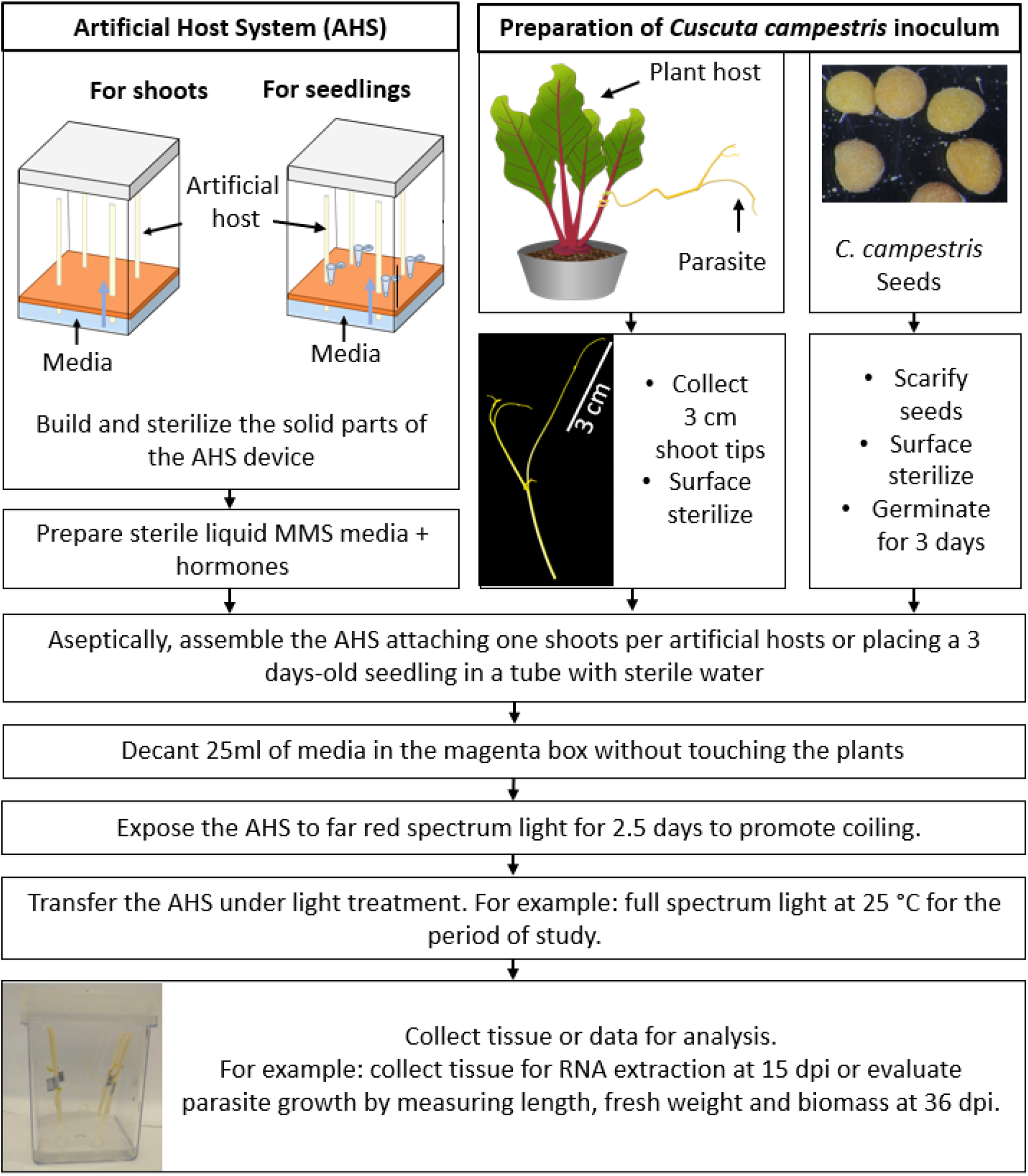
Schematic depiction of the artificial host system (AHS) and the key steps needed to grow *Cuscuta campestris* on the artificial host. Days post inoculation (dpi).

### Media preparation

Modified Murashige and Skoog liquid media (MMS) (Srivastava & Dwivedi, 2001) was used as base media for most of the experiments. It contained 300 mg l^-1^ NH_4_NO_3_, 800 mg l^-1^ KNO_3_, 250 mg l^-1^ CaCl_2_.2H_2_O, 260 mg l^-1^ MgSO_4_.7H_2_O, 120 mg l^-1^ KH_2_PO_4_, and 3% glucose, pH 5.7. Sterile 1X MS micronutrients (Phytotechlab, USA) were added after autoclaving. This MMS media was supplemented with filter-sterilized phytohormones 1-naphthaleneacetic acid (NAA) (3 mg l^-1^) or 6-benzylaminopurine (BA) (1 mg l^-1^) for specific experiments.

### Preparation of parasitic plant material

Vegetative *C. campestris* shoot tips were used as inoculum for most of the experiments. Nurseries of the parasite were established on beets (*Beta vulgaris,* Detroit Dark Red) as follows. *C. campestris* seeds were scarified by immersion in sulfuric acid (95%) for 20 min, followed for 3 washes with water for 10 minutes. Seeds were germinated on wet filter paper in Petri dishes for 4 days, then seedlings were attached to petioles of one-month-old beet plants using a small piece of tape. Host and parasite were placed under direct full spectrum light (Mercury-Vapor lamp) and far-red enriched light (Sylvania Spot-Gro bulb, enriched in >750nm wavelength, 35 µmol m^-2^s^-1^) under 14h light /10h dark photoperiod. Plants were watered as needed and fertilized with All Purpose Plant Food (Miracle-Gro, USA). Once the *C. campestris* plants produced large numbers of shoots (around 1.5 months after inoculation), 3 cm shoot tips were harvested for inoculum. Shoot tips were surface sterilized with 3.5% sodium hypochlorite and 0.04% Tween20 for 10 min, followed by 5 washes with sterile water just before attaching them to the artificial host.

When seedlings were used as inoculum, parasite seeds were scarified as described above and surface sterilized for 5 minutes with 3.5% sodium hypochlorite and 0.1% SDS followed by 5 washes with sterile water. Sterilized seeds were germinated as described above under sterile conditions. For the germination of seeds collected from plants growing in the AHS, mature seeds were scarified and germinated as described above, except the scarification period was 7 min. Seedlings and new shoots were attached to *Arabidopsis thaliana* inflorescences under light conditions described above. Aracones (Arasystems, Belgium) were used to increase relative humidity in the parasite-host microenvironment and to avoid the parasite growing onto neighboring plants in later stages.

### Artificial host system assembly

Working under aseptic conditions, surface-sterilized *C. campestris* shoots were attached to a spindle using a sterile strip of aluminum foil that held the parasite shoot securely to the spindle with a gentle twist of the foil (e.g., Fig. **2a,b**). The foil was placed approximately 1 cm from the tip of the *C. campestris* shoot, and each parasite shoot was placed high enough on the spindle to keep its lowest extremity away from any contact with media in the bottom of the box. After attaching the shoots, liquid media was added to the box, avoiding touching the plants. Magenta boxes were sealed with parafilm and placed under a Sylvania Spot-Gro bulb for 2.5 days with a 14h light/10h dark photoperiod at 33°C. Boxes were then transferred to a growth chamber under full spectrum light (61 µmol m^-2^s^-1^, 25°C). Relative humidity in the boxes with liquid media reached 100%, whereas boxes without media were at 43%. AHS boxes were kept under full spectrum light at 25°C for 33.5 days in the case of growth assays, this time was reduced to 12.5 days for other assays like RNA-seq. When seedlings were used as inoculum, the procedure was the same except a 3-day-old *C. campestris* seedling was placed in a 0.2ml microcentrifuge tube containing 150 µl of sterile water to keep the seedling hydrated and the tube was attached to the base of each spindle.

**Fig. 2.**
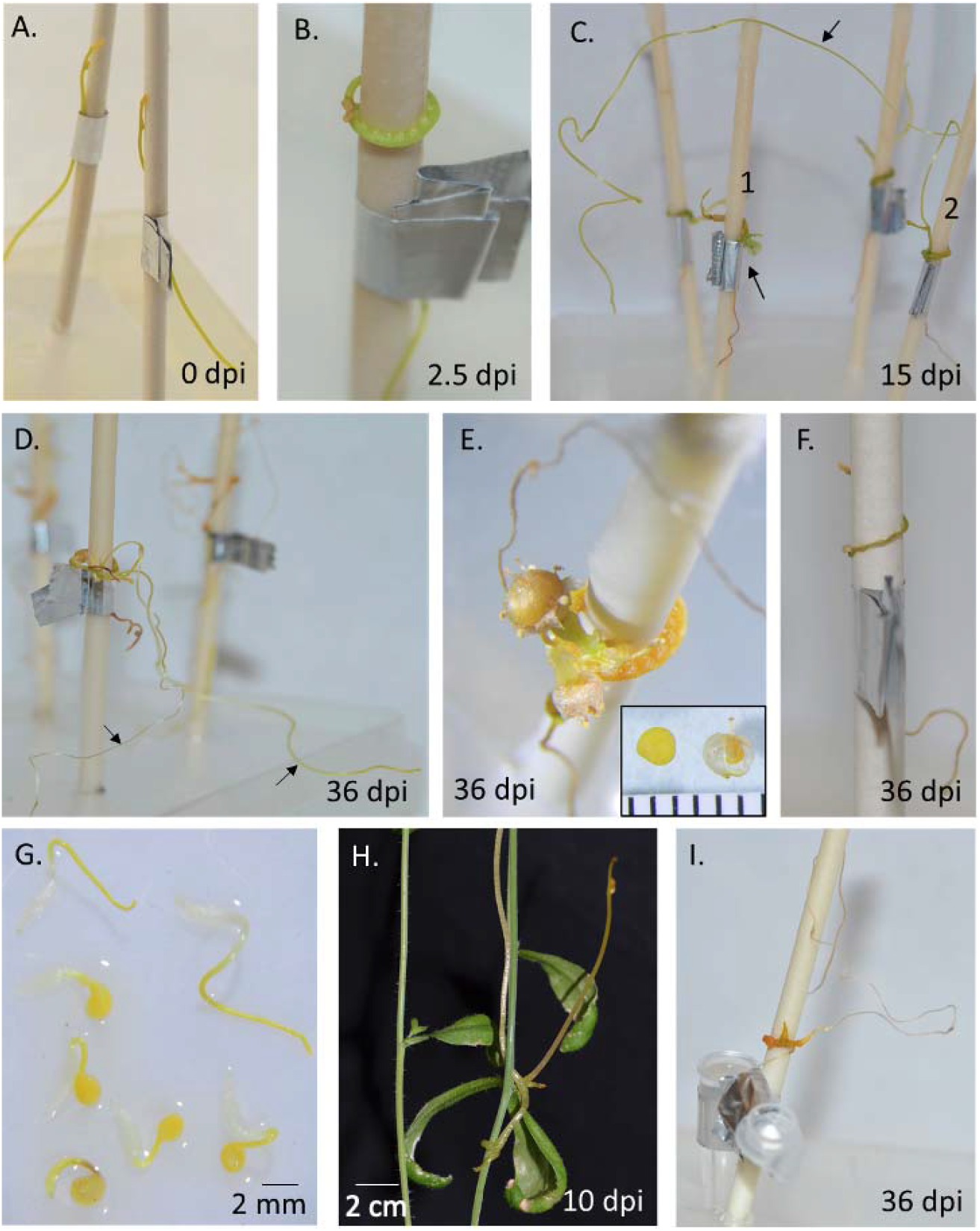
*Cuscuta campestris* grows without a plant host under the controlled conditions of the artificial host system (AHS). a-f. *C. campestris* 3 cm shoot tips were used as inoculum. a. 3 cm shoot tips attached to an artificial host at 0 days post inoculation (dpi). b. Parasite coiled with holdfasts after 2.5 dpi. c. Parasite at 15 dpi, growth is evident as flower development (arrow) in plant No. 1 and production of a new shoot in plant No. 2 d. New shoots elongated, 36 dpi. e. Parasite flowered and produced seeds (inset; scale is in millimeters). f. *C. campestris* attached to an artificial host without any media or water at 36 dpi. Parasite did not develop swollen holdfast or subsequent growth. g. Seedlings after scarification and germination of seeds obtained from *C. campestris* grown in the AHS. h. Seedling from seed obtained through AHS colonizing *A. thaliana* 10 dpi. i. Seedling growing 36 dpi after being used as inoculum in a second AHS.

### Measuring parasite growth

For evaluating parasite growth, AHS boxes were opened, plants were detached from artificial hosts, extended on a white surface with a ruler, and photographed. Pictures were analyzed using ImageJ 1.51j8 (National Institutes of Health, USA) to obtain the total length of the plant, including all branches and the initial 3 cm shoot. Shoots were then weighed to obtain fresh weight and dried at 50°C for four days to before reweighing to obtain biomass data (dry weight). Data were collected from three independent experiments with a total of 96, 105 and 76 plants, respectively. Only plants that coiled and produced healthy haustoria were considered for growth analyses. Statistical analysis used JMP Pro 15 (SAS Institute Inc.).

### CFDA Assay

*C. campestris* plants were grown for 15 days in AHS containing MMS media with NAA and BA. The spindle “hosts”, along with the connected parasites, were then transferred to a 1.5-ml microcentrifuge tube containing the same media plus 0.3 mg ml^-1^carboxyfluorescein diacetate (CFDA, initially dissolved in DMSO) and held inside a closed sterile glass tube. Negative controls consisted of the equivalent concentration of DMSO without CFDA. In this way, the spindle conducted media containing CFDA to the point of the parasite attachment that was the only point of transfer to the parasite. After four days, plants were detached, rinsed thoroughly with water, and examined for CFDA fluorescence using a stereoscope under UV light with an EGFP filter.

### Microscopy

*C. campestris* was grown in the AHS with MMS, NAA and BA. Plants were collected 16 days post inoculation (dpi), hand-sectioned, and stained with Toluidine Blue-O. Cross-sections were evaluated under an inverted microscope (Olympus CKX53) equipped with a camera (Olympus EP50).

### RNA-seq and transcriptomic analysis

Vegetative parasite shoot tips were introduced to the AHS containing MMS media with NAA and BA. After 15 dpi, tissue was collected for RNA extraction (Fig. **S4**). Plants with healthy holdfasts and newly developed shoots were detached from the artificial host. For haustorial samples, regions harboring visibly healthy holdfasts (identified by formation of new shoots) were separated from any shoot or lateral meristem. For non-haustorial stem samples, new healthy and turgid shoots were collected after removing 5 mm of the stem connecting to the haustorial region to avoid tissue contamination. All tissues were immediately frozen in liquid nitrogen. Samples from 3-4 plants were pooled for each replicate and four replicates were used for each type of tissue. For RNA extraction, frozen tissue was ground using glass beads in a bead beater and extracted using the RNeasy Plant Mini Kit (Qiagen). RNA was precipitated using 3M NaOAc and Ethanol 100%, and resuspended in RNAse-free water. RNA concentration and purity were evaluated using a Nanodrop OneC (ThermoScientific), and quality was confirmed using an RNA TapeStation System (Agilent). A mRNA library was prepared using Poly-A enrichment, and sequencing was performed by Novogene using Illumina NovaSeq 6000 platform with PE150.

### Bioinformatic analysis

RNA-seq raw data quality was initially evaluated using FASTQC v.0.11.9 (Braham Bioinformatics). Then, FASTP v.1.0.0 (Chen *et al.*, 2018) was used to detect and trim adaptors, and filter out sequences shorter than 50nt, with low quality or low complexity. Clean reads were mapped with STAR 2.5.2b (Dobin *et al.*, 2013) against the genome of *C. campestris* (Vogel *et al.*, 2018). Gene counts obtained using STAR 2.5.2b were processed by DESeq2 1.30.1 (Love *et al.*, 2014) using R (https://www.r-project.org/), with differential expression analysis using the default settings and a threshold of genes with more than 10 counts. To increase the stringency of differentially expressed gene (DEG) detection and filter out biases related to the structured distribution of the samples (haustoria vs stem from the same plant pool), gene counts were also processed by edgeR 3.32.1 (Robinson *et al.*, 2010; McCarthy *et al.*, 2012) using R to perform a second DEG analyses accounting for plant pool as blocking factor using the option of generalized linear model - Likelihood Ratio Test (glmLRT). Genes with low counts were filtered using count-per-million (CPM) and normalization was done using trimmed mean of M-values (TMM) according to default settings. DEGs were determined by the genes in the intersection from the two approaches and considering thresholds at adjusted p-value (padj) < 0.05 for DESeq2 or False Discovery Rate (FDR) < 0.05 for edgeR.

Protein sequences reported for *C. campestris* (Vogel *et al.*, 2018) were used for a functional annotation using eggNOG-mapper v2 (Huerta-Cepas *et al.*, 2017, 2019) to obtain their associated GO terms. Then a Kolmogorov-Smirnov like enrichment test (also known as GSEA) was performed. Enrichment analysis was done independently for down- and up-regulated genes. DEGs, their expression p-value according to DESeq2, and their GO terms were analyzed by topGO R package version 2.42.0 (Alexa & Rahnenfuhrer, 2021) considering the settings weight 01 and NodeSize 10. Top enriched GO terms (p-value ≤ 0.001) were plotted using REVIGO (Supek *et al.*, 2011) under default conditions.

### Comparison with other *C. campestris* expression datasets

Genes found to be up-regulated in the haustorial region of *C. campestris* growing in AHS (with a padj < 0.01) were compared to other published gene expression datasets (Fig. **S4**). The first selected dataset corresponds to up-regulated genes in micro-dissected haustoria tissue, 87 hours after the forced induction of haustoria while in contact with a leaf of *A. thaliana* (Kaga *et al.*, 2020). The second dataset corresponds to up-regulated genes in fully formed haustorial region of *C. campestris* (formerly published as *C. pentagona*) growing on two hosts, *Solanum lycopersicum* and *Nicotiana tabacum* (Ranjan *et al.,* 2014). A third dataset was taken from Kim, *et al.* (2014), that used functional connections between *C. campestris* (also formerly published as *C. pentagona*) and *A. thaliana*. In this case, we used clean gene counts of total expressed genes in the interface (corresponding to the haustorial region) and parasite (stem tissue) to perform DEG analyses. Data corresponding to three replicates was processed by to edgeR 3.32.1 1 (Robinson *et al.*, 2010; McCarthy *et al.*, 2012) to perform a DEG analysis accounting for batch effect and using default settings. DEGs were determined by FDR < 0.05.

Genes IDs used by Vogel *et al.* (2018) for *C. campestris* were used as a convention. A BlastX (−evalue 1e-50, −max_target_seqs 1, −max_hsps 1) against the proteins reported by Vogel *et al.* (2018) was used to retrieve the gene IDs that correspond to the data obtained from Ranjan *et al.* (2014) and Kim *et al.* (2014). Gene IDs utilized by Kaga *et al.* (2020) were used directly as they correspond to Vogel *et al.* (2018) gene IDs. We grouped the datasets of Ranjan *et al.* (2014) and Kim, *et al.* (2014) based on their study of functional haustorial regions of *C. campestris* growing on a plant host.

Once the shared upregulated genes in haustorial regions among the selected studies were identified, we used the *C. campestris* protein sequences in Vogel *et al.* (2018) to identify their homologues in *A. thaliana* TAIR10 (Phytozome v12.1.6) using BlastX (−evalue 1e-50, −max_target_seqs 1, −max_hsps 1). Candidate *A. thaliana* homologues were subjected to Single Enrichment Analysis (SEA) using agriGO v2.0 (Tian *et al.*, 2017) using Go slim and Phytozome v11 as reference, with default settings.

## Results

### An artificial host system for growing *C. campestris* under axenic conditions

We developed a system that provides the essential components for *C. campestris* growth in a way that mimics growth on a plant host. The main elements of the AHS are a solid vertical support with capillary capacity, liquid media with proper nutrients and phytohormones, and a translucent container to hold these together under aseptic conditions (Fig. **1**, **S1**). We evaluated several materials for use as artificial hosts, and found the best to be the paper spindle from a cotton swab, which has a firm structure, yet sufficient porosity to allow the movement of liquid media by capillary action. This vertical artificial host “stem” provides a tactile stimulus that, when combined with far-red enriched light, induces the parasite to coil and form haustoria (Tada *et al.*, 1996; Furuhashi *et al.*, 2011). The AHS works with *C. campestris* shoots originating from either vegetative cuttings or seedlings (Fig. **1**). However, shoot tips exhibited a higher rate of coiling and holdfast production (84% and 82%, respectively) than seedlings (74% and 49%, respectively). Therefore, we used shoot tips for further experiments.

### *C. campestris* displays all lifecycle stages under the AHS conditions

*Cuscuta* species are obligate parasites, so normally cannot grow beyond the seedling stage without a plant host (Kaiser *et al.*, 2015), but the AHS adequately substituted for the host, enabling *C. campestris* to complete all life stages. We attached 3-cm shoot tips to artificial hosts and evaluated their development up to 36 dpi. At 2.5 dpi, parasites had coiled and formed holdfast structures on the shoot in contact with the artificial host (Fig. **2a,b**). We consider the presence of a holdfast to be a sign of initiation of haustorial development. At 15 and 36 dpi parasites had conspicuous enlargements in the haustorial region and produced new vegetative shoot growth, flowers, or both (Fig. **2c-e**). After 36 dpi, some plants were still elongating (Fig. **2d**, **S3a**) whereas others had long shoots that had died back to a small section (< 3 cm) of green turgid tissue at the tip, which we considered to be senescent shoots (Fig. **S3a**). In the cases where artificial hosts were supplied with pure water or no media at all, *C. campestris* was not able to grow (Fig. **2f**, **S2**). Generally, no healthy holdfasts developed under these treatments, but some formed when just water was applied, resulting in a slight increase in fresh weight. However, addition of nutrient media led to significant gains in length and biomass (Fig. **S2**).

*C. campestris* colonizes new hosts through either vegetative shoot branches or seedlings coming from germinated seeds. To be useful, the AHS should be able to support growth of *C. campestris* from both sources. We confirmed vegetative propagation using shoot cuttings from AHS-grown plants at 36 dpi by taking active shoots of at least 3 cm and applying them to new plant hosts (Fig. **S3**) as well as new artificial hosts. These shoots were viable, coiled, produced holdfasts, and grew on both types of hosts. For propagation by seed, we scarified the AHS-produced seeds and determined that they are able to germinate (Fig. **2g**), with a germination rate of 28% (n=47). We attached these seedlings to new plant hosts as well as new artificial hosts and found that they are able to grow on plant hosts as well as on artificial hosts (Fig. **2h,i**).

### The parasite in the AHS feeds through the haustorial region

The haustorium is the signature organ of parasitism in parasitic plants and is essential for uptake of water and nutrients from the hosts (Kuijt, 1977). To characterize the ability of haustoria to take up resources, we added the phloem-mobile dye CFDA (Grignon *et al.*, 1989) to the media and monitored its uptake and translocation into the parasite. CFDA is not fluorescent, but once inside a cell it is cleaved to carboxyfluorescein, which is fluorescent. We observed the assimilation of fluorescent signal in the parasite tissues, both in association with the holdfast/haustorium region and in scale leaves that cover apical and lateral meristems, which are located away from the haustorial region (Fig. **3**). These observations indicate that the haustoria of *C. campestris* growing on the AHS function in uptake of small molecules.

**Fig. 3.**
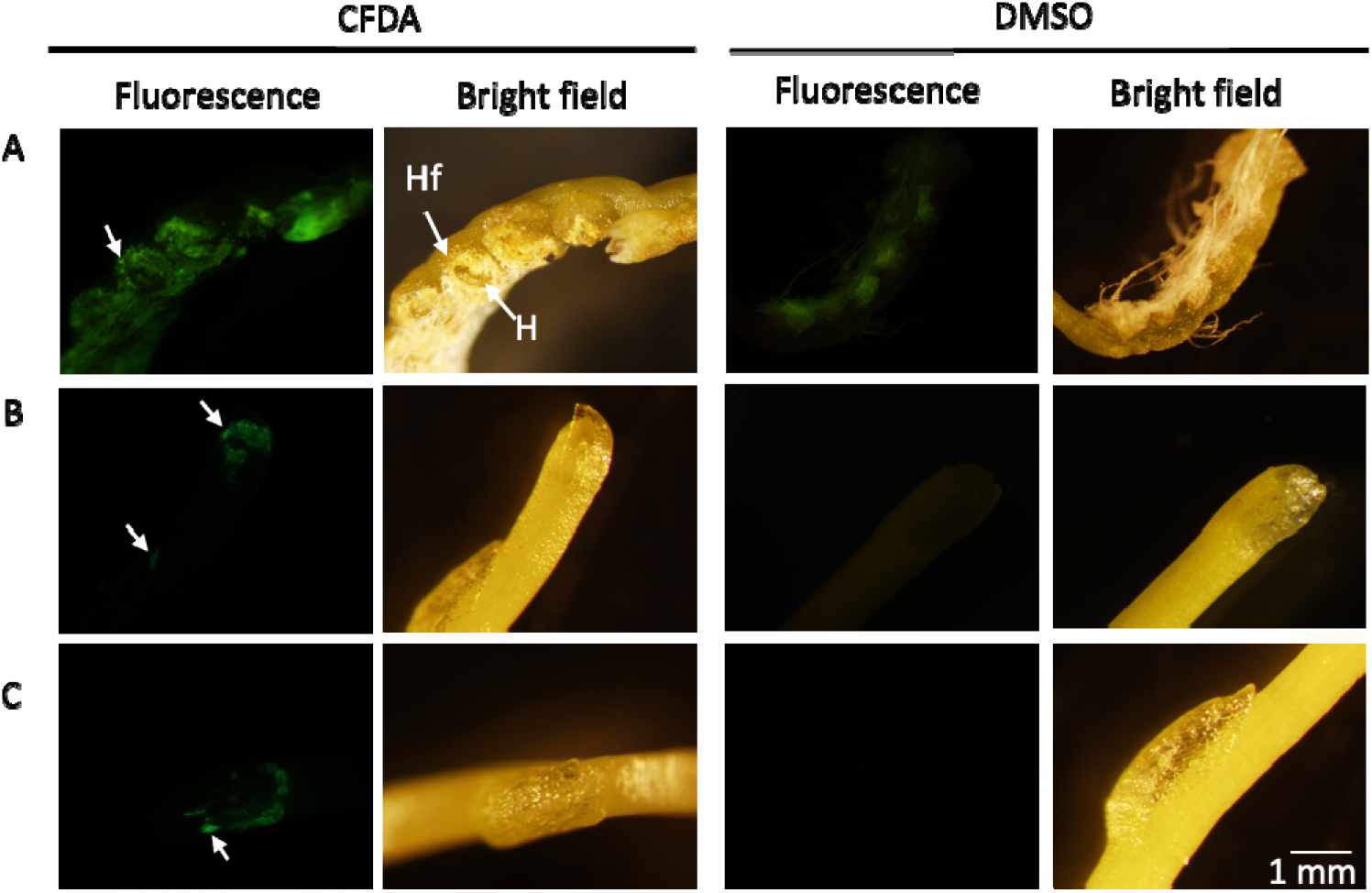
*Cuscuta campestris* growing in artificial host system (AHS) exposed to the fluorescent dye carboxyfluorescein diacetate (CFDA) or DMSO solvent control from the artificial host. a.Haustorial region. Cells of the holdfast show fluorescence. Holdfast (Hf), Haustorium (H). band cScale leaves protecting apical and lateral meristems.

### The AHS allows study of *C. campestris* under controlled axenic conditions

#### Establishment: Parasite establishment is independent of the media composition, although holdfast morphology is sensitive to external phytohormones

To evaluate the impact of media composition on parasite establishment (here defined as stem coiling and formation of the holdfast), we applied *C. campestris* shoot tips to artificial hosts supplied with six different media treatments: nothing (no media at all), pure water, MMS media, MMS with NAA, MMS with the BA, and MMS with NAA plus BA. The coiling phase occurred by 1 dpi and we found no significant differences in the coiling rates among the treatments (Fig. **4a**). The holdfast formation phase occurred by 2 dpi, and the percentage of formation was similar among all treatments except for the dry spindle (Fig. **4g**). The media composition did affect holdfast morphology. At 8 dpi, holdfasts showed altered phenotypes in response to the different media compositions (Fig. **4b-g**). Parasite holdfasts exposed to MMS media with NAA and or BA exhibited a more complex structure than ones exposed to MMS media or water, and inclusion of BA produced the greatest effect, leading to more developed and thicker holdfasts (Fig. **4c,e**) that more closely resembled those formed on a plant host such as *N. benthamiana* (Fig. **4h**). To further examine the parasitic organ produced on the artificial host, we inspected the anatomy of the haustorial region. The holdfast cells form an adhesion disc on the outer section of the haustoria that adhered firmly to the artificial host and resisted removal as evidenced by spindle fibers remaining attached to the cells (Fig. **5a-c**). Cross-sections of the holdfasts revealed that each haustorial region contains a range of holdfasts at different developmental stages (Fig. **5d-f**). Immature organs, usually located at the extremes of the haustorial region, had holdfasts but no haustorial development (Fig. **5d**). Mature organs with a developed haustorium were often located in the center of the haustorial region, and these showed substantial protrusion toward the artificial host (Fig. **5e,f**). Within the files of elongated cells radiating toward the host, tracheary elements of xylem were identifiable by their ringed secondary walls (Fig. **5e,f**). Although our sectioning did not reveal a continuous file of tracheary elements, we observed them both near the parasite stem and close to the distal part of the haustorium that contacts the host surface (Fig. **5e,f**).

**Fig. 4.**
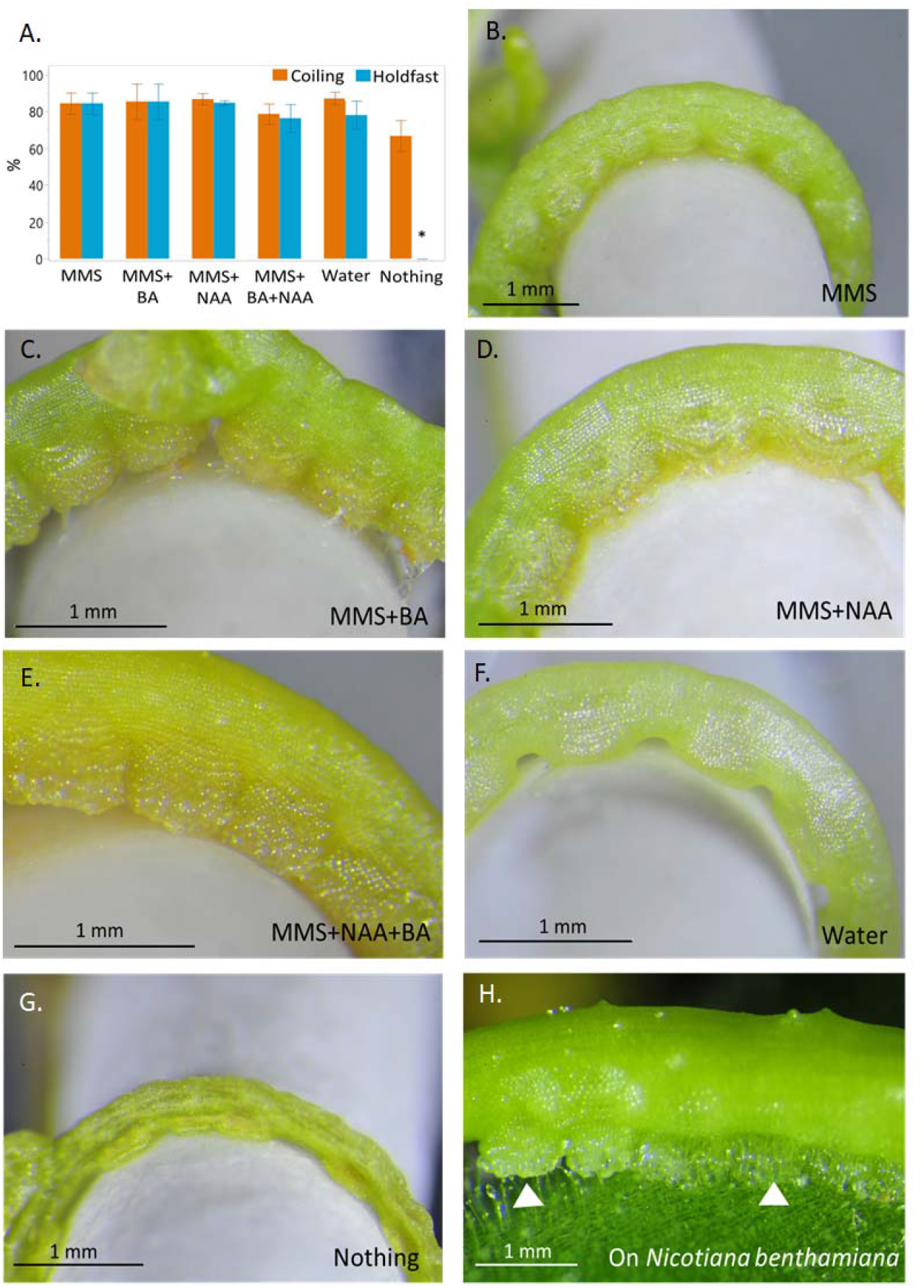
Characterization of *Cuscuta campestris* coiling and holdfast morphology in the artificial host system (AHS). a. Frequencies of coiling and holdfast production evaluated at 36 days post inoculation (dpi). The experiment was repeated three independent times, with an average of 15 replicates. Graph presents means between the three experiments and ±SE. b-g. Holdfast morphology as influenced by phytohormones in the media. Holdfasts of parasites growing with MMS (Modified Murashige and Skoog) media (b) c.MMS+6-benzylaminopurine (BA) (c), MMS+1-naphthaleneacetic acid (NAA) (d), MMS+NAA+BA (e) or just water (f). g. Parasite growing on an artificial host without any media, “Nothing”. h. Holdfasts (white arrows) of *C. campestris* growing on *Nicotiana benthamian*a. Pictures were taken at 8 dpi.

**Fig. 5.**
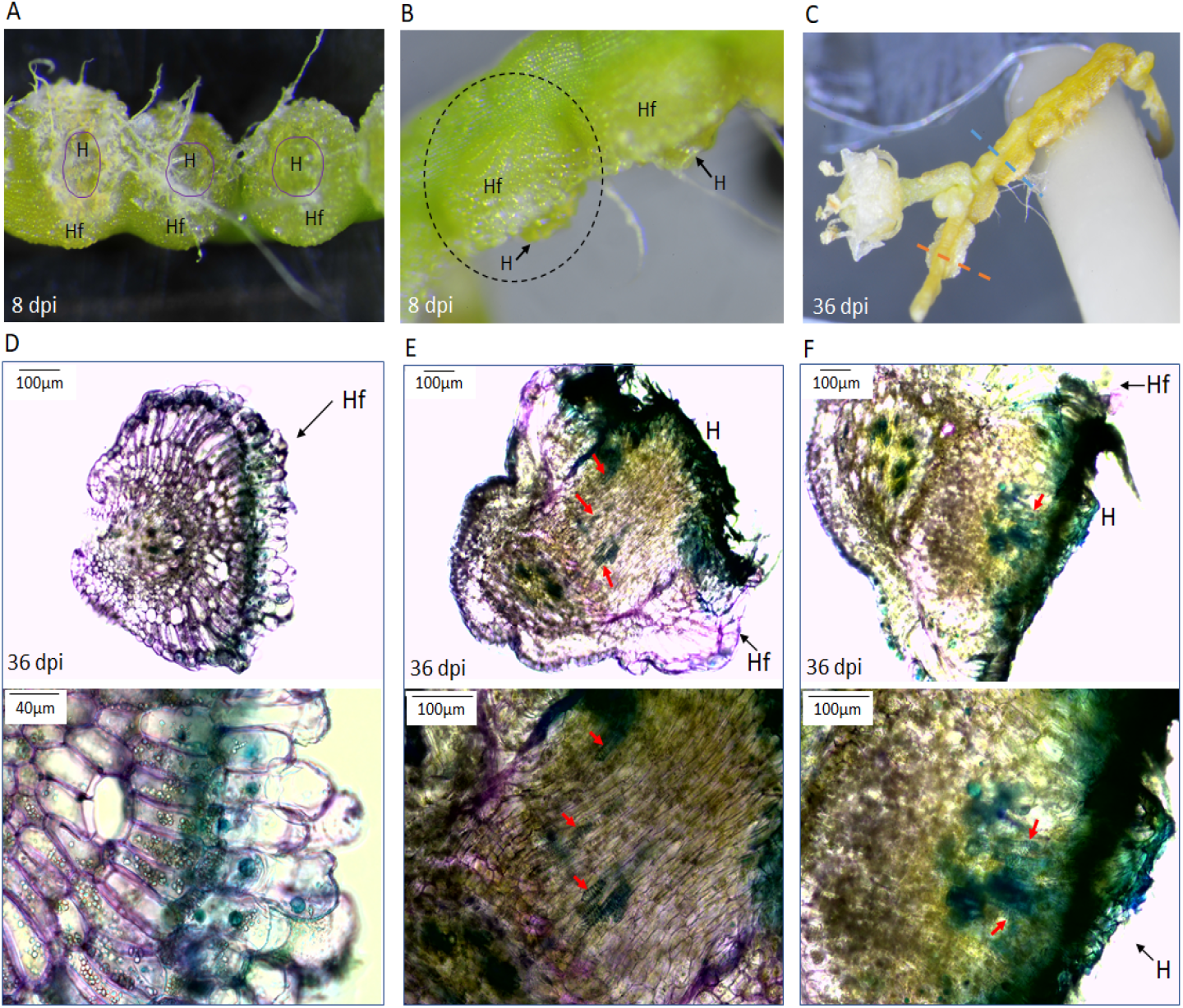
Anatomy of parasitic organs produced by *Cuscuta campestris* growing on an artificial host with MMS (Modified Murashige and Skoog) media including 1-naphthaleneacetic acid (NAA) and 6-benzylaminopurine (BA). a. haustorial region inner face view at 8 days post inoculation (dpi). Haustoria (H) is observed in the center of the parasitic organ surrounded by the holdfast (Hf). Paper fibers from the artificial host are visible. b. Lateral view of haustorial region with visible holdfast and protruding haustoria at 8 dpi. Dotted line circle indicates a parasitic organ. c. haustorial region with parasitic organs at different developmental stages (36 dpi). d-f. Cross sections of parasitic organs of *C. campestris* growing on an artificial host at 36 dpi. Bottom panels show higher magnification of upper panels. d. Parasitic organ in an early developmental stage corresponding to the orange dotted line in Fig. c. Holdfast epidermal cells on the face were in contact with the host, so represent the adhesion disk. Higher magnification (bottom panel) shows these cells to be elongated. eand f. Mature parasitic organs corresponded to the blue dotted line in Fig. c. Haustorium development is conspicuous with elongated cells protruding towards the artificial host. Hand-cut cross-sections were stained with Toluidine blue-O to reveal tracheary elements of the xylem (in blue, indicated by red arrows).

#### Parasite growth and development: Phytohormones affect *C. campestris* weight, length, and fitness

To evaluate the effect of the AHS media composition on parasite growth, we inoculated *C. campestris* shoot tips on artificial hosts and evaluated changes in parasite fresh weight, biomass, and total length at 36 dpi. The parasite exhibited significantly higher fresh weight and biomass when exposed to MMS media containing phytohormones as compared to only MMS media or water (Fig. **6a,b**). The greatest gain in fresh weight and biomass occurred in treatments containing BA compared to MMS alone or containing only NAA. Parasites exposed to only MMS or water treatments showed comparable fresh weights, while MMS allowed significantly higher biomass accumulation than just water. Fewer differences were found when measuring shoot length. Parasites exposed to media with MMS and BA attained the greatest length but did not differ significantly from plants with NAA-containing media. No significant length gain was found in parasites growing on AHS media without phytohormones (Fig. **6c**). Our results show that the fresh weight, biomass, and total length of *C. campestris* are significantly enhanced by externally-provided phytohormones.

**Fig. 6.**
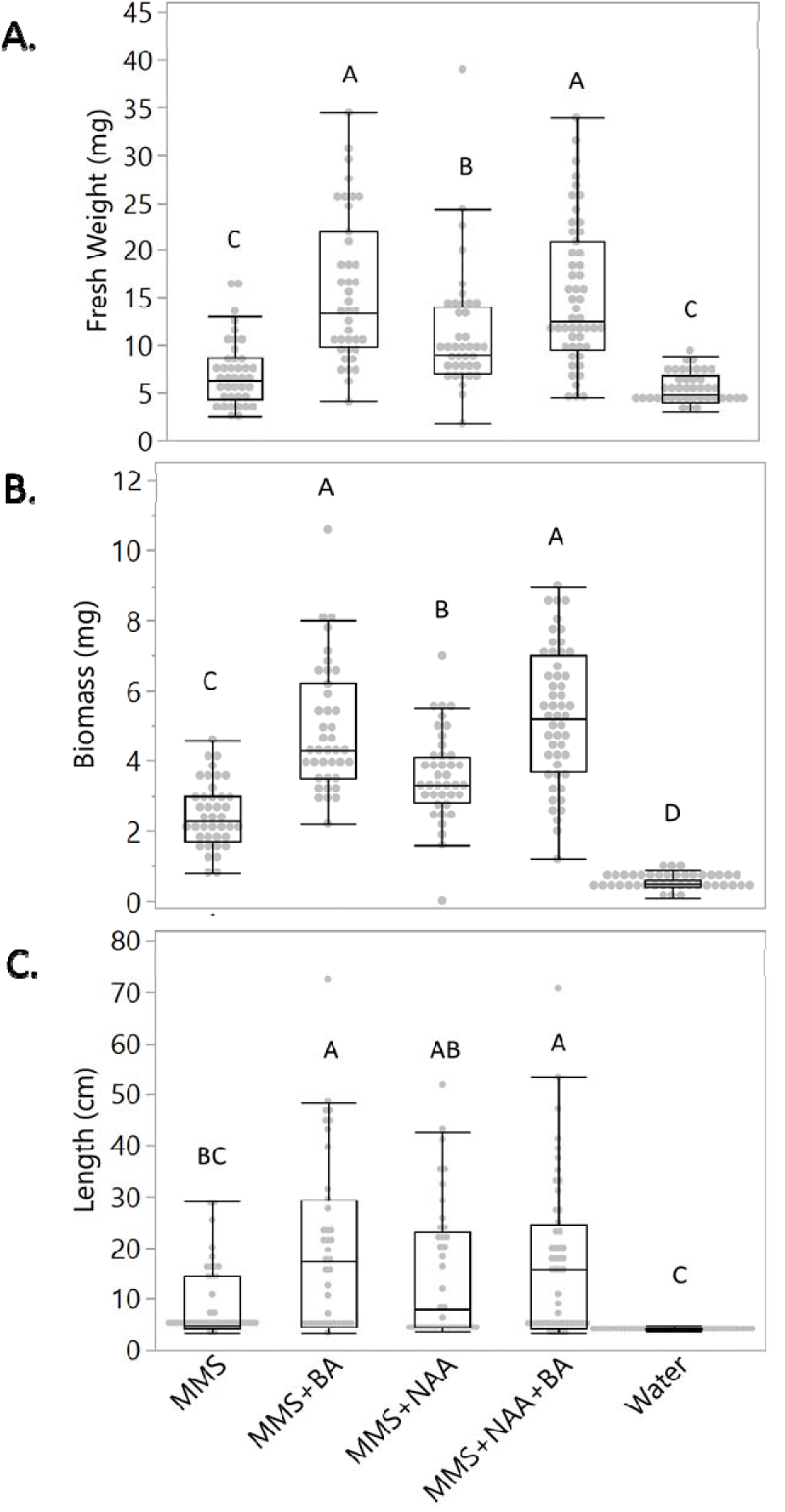
*Cuscuta campestris* growth in the artificial host system (AHS) supplied with different media. Parasite fresh weight (a), parasite biomass (b) and parasite total length (c) were measured at 36 days post inoculation. Different media were tested: MMS (Modified Murashige and Skoog) media, MMS with 1-naphthaleneacetic acid (NAA), MMS with 6-benzylaminopurine (BA), MMS with NAA and BA, and water alone. Statistical differences were detected by ANOVA with post-hoc Tukey HSD test. Differences were considered statistically significant at P<0.05 and indicated with different letters. Data are pooled from three independent experiments. Analysis included only plants that coiled and developed healthy holdfasts. Number of samples ranged from 39 to 51 per treatment.

*C. campestris* propagation occurs through seeds and shoots, so measuring these aspects of parasite growth in the AHS provides an indication of *C. campestris* fitness. We assessed the percentage of new active shoots and fruits as indicators of the parasite potential to propagate and colonize new hosts. We consider actively growing shoots to be those with turgid shoot tips of at least 3 cm and able to colonize new hosts (Fig. **S3**). Only plants growing with media containing BA showed a significant increase in the percentage of active shoots (Fig. **7a**). With respect to sexual reproduction, flowers were formed under all MMS media conditions, regardless of phytohormones (**Fig. 7b**). Parasites with fruits generally produced one fruit with 1 to 2 viable seeds. Only *C. campestris* exposed to media containing phytohormones produced seed. These results suggest that externally supplied phytohormones BA and NAA enhance parasite fitness in the AHS.

**Fig. 7.**
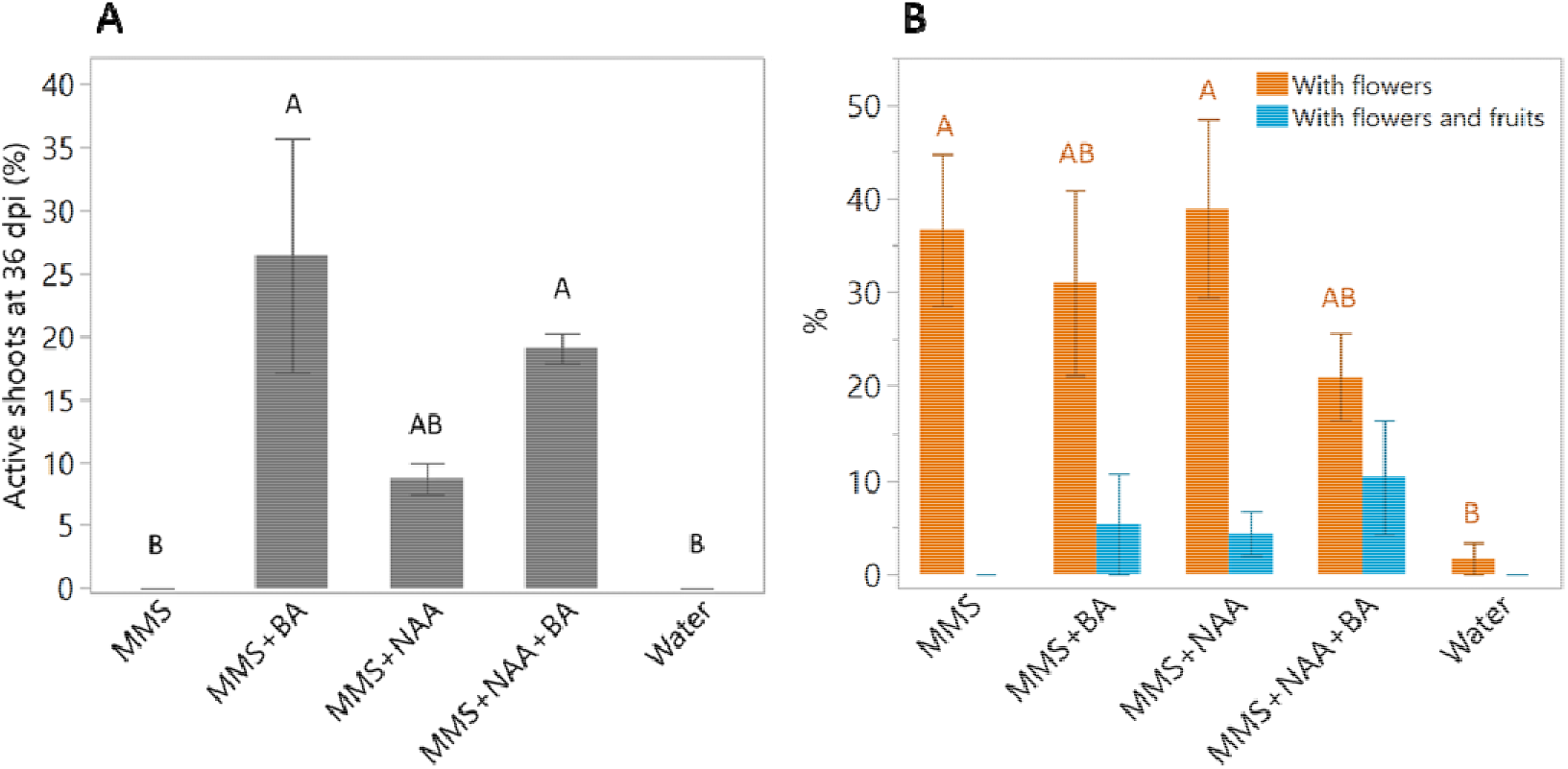
Effect of artificial host system (AHS) media composition on the ability of *Cuscuta campestris* to generate parasitically competent shoots, flowers, and fruits. a. The percentage of plants with active shoots after 36 days post inoculation (dpi). Active shoots were defined as those with tips of at least 3 cm of healthy, turgid shoot, which are characteristic of shoots capable of forming new host connections. b. The percentage of plants with flowers only or with fruit production. Analyses included only plants that coiled and developed a healthy holdfast. Graph presents means ±SE between three independent experiments, each experiment with a number of plants ranged from 6 to 19 per treatment. Statistical differences were detected by ANOVA with post-hoc Tukey HSD test. Differences were considered statistically significant at P<0.05 and indicated with different letters.

In measuring *C. campestris* fitness and growth, we noted a frequent tradeoff in type of growth. Some plants grew long vegetative shoots, while others would thicken and produce flowers. In order to understand parasite growth parameters, we evaluated the relationship between biomass, fresh weight, and total length. Data from each parameter were grouped, regardless of the media, and Pearson’s coefficient of correlation was calculated between the pairs of parameters (Fig. **S4**). We found the strongest correlation between biomass and fresh weight (r=0.869). We also analyzed the distribution of plants with flowers and fruits in relation to biomass and total length, and this confirmed that most parasites with flowers or flowers and fruits displayed reduced total length and elevated biomass compared to the parasites without flowers or fruits (Fig. **S5**). We conclude that reliance on any single parameter can be misleading and recommend the use of more than one of these three growth parameters to capture the phenotypic response of *C. campestris*.

#### Haustorial function: Characterization of the gene expression profile of a functional haustorial region in the absence of a plant host

In order to explore features associated with a functional haustoria in the absence of a plant host, we analyzed differential gene expression between the haustorial region and the newly developed stem of plants growing in the AHS (Fig. **S6**). We grew *C. campestris* in the AHS (MMS media with NAA and BA at 15 dpi) and sequenced mRNA extracted from the haustorial region and vegetative shoots. On average, 90% of the reads uniquely mapped to the previously sequenced *C. campestris* genome (Vogel *et al.*, 2018). We found a total of 20,749 DEG, with 9,939 up-regulated and 10,810 down-regulated in the haustorial region compared to stem (Fig. **S6**).

We performed a Gene Ontology (GO) enrichment analysis to determine which biological processes were associated with AHS grown haustorial development and function. Among the overrepresented processes in the haustorial region are lignin biosynthesis, xylem development, and response to hormones, which may be related to haustoria development (Fig. **8a**, Table S1). Other enriched processes include transport (ions, carbohydrates and amino acids), hormones biosynthesis, response to abiotic stress, and defense, which may be associated with haustorium interaction with a host. On the other hand, genes underrepresented in the haustorial region compared to the shoot include those functioning in organ and stomatal morphogenesis, gravitropism, response to hormones, cuticle and cell wall-related biosynthetic processes, auxin efflux, cell differentiation, DNA-related processes, and translation (Fig. **8b**, Table S1).

**Fig. 8.**
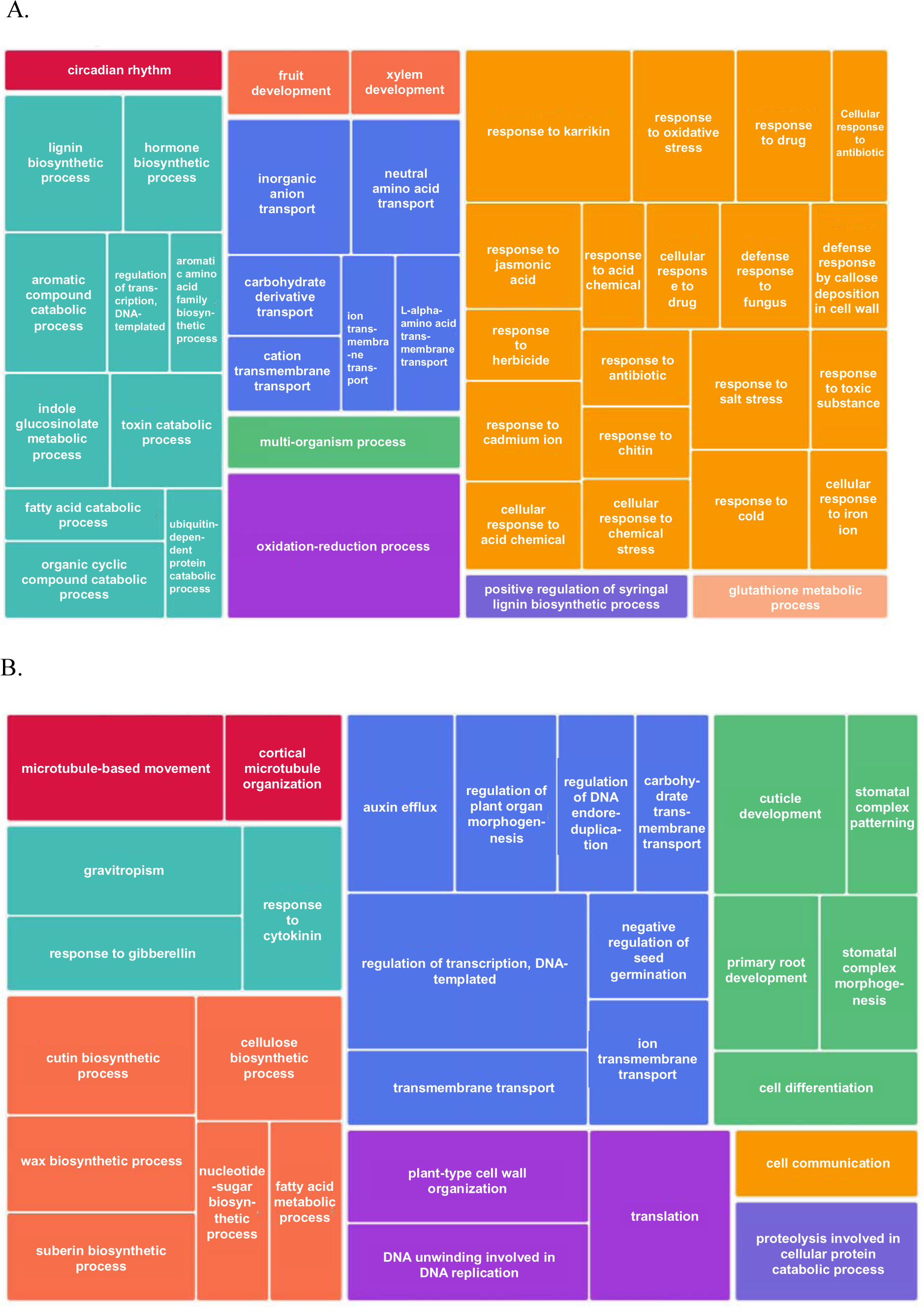
Enriched Gene Ontology (GO) terms associated to Biological Process corresponding to differentially expressed genes (DEGs) in haustorial region of a parasite growing in artificial host system (AHS). a. 9,939 up regulated genes were included. After the enrichment analysis, the top 55 enriched GO Terms (enrichment p-value < 0.001) were selected to build a TreeMap. b. 10,810 down regulated genes were included. After the enrichment analysis, the top 35 enriched GO Terms (enrichment p-value < 0.001) were selected to build a TreeMap. Kolmogorov-Smirnov like enrichment tests (also known as Gene Set Enrichment Analysis - GSEA) was performed for aand b, considering expression p-value. Each rectangle is a single cluster representative, its size represents the frequency of the GO term in the underlying GO Annotation database. Colors indicate superclusters of loosely related terms.

#### Genes upregulated in haustorial regions of plants in the AHS are also expressed during parasitism of actual plant hosts

Finally, we compared the DEG found in the haustorial region of a parasite growing in AHS with other published gene expression datasets from haustorial tissues associated with real plant hosts. We used datasets from two categories of haustoria, “functional” or “immature”. Functional haustoria come from studies that report established haustoria that allow parasite growth. Data in the “functional” category was derived from a union of data from analysis of *C. campestris* growing on *A. thaliana* inflorescences (Kim *et al.*, 2014) and on *S. lycopersicum* and *N. tabacum* stems (Ranjan et al., 2014). Immature haustoria have visible holdfasts, but growth of new shoots or flowers is not reported. Data in the “immature” category are from *C. campestris* haustoria 87 hours after haustorial induction in contact with a leaf of *A. thaliana* (Kaga *et al.*, 2020).

Comparison of induced DEGs from the AHS-grown haustoria to the functional and immature haustoria in contact with a plant host data revealed 220 genes shared by all three datasets regardless of haustorial functional status or contact with actual plant tissues (Fig. **S7**) (Table **S2**). Another 62 genes were uniquely shared by functional haustorial tissues even though AHS-grown plants lacked contact with a real plant host. To assess functions of the 220 and 62 shared genes, we searched for homologues of the *C. campestris* genes in *A. thaliana,* finding 148 and 45 genes, respectively (Fig. **S8**). Genes present in all haustoria were grouped in biological processes categories related to anatomical structure development, multicellular organism development, and response to endogenous stimulus, biotic stimulus, external stimulus, abiotic stimulus, and stress (Fig. **S8a**). Genes shared by just functional haustoria were grouped in biological processes related to anatomical structure development, response to abiotic stimulus and response to stress (Fig. **S8b**).

## Discussion

### Features of the AHS

One of the primary challenges of studying obligate parasitic plants is the difficulty in separating the biology of the parasite from that of its host. Although *Cuscuta* species have previously been grown under tissue culture conditions (Loo, 1946; Baldev, 1962; Bakos A. *et al.*, 1995; Srivastava & Dwivedi, 2001; Borsics & Lados, 2002; Švubová & Blehová, 2013), the phenotype of the parasite under such conditions does not correspond to the coiling and haustorial feeding habit that characterizes normal growth. To overcome this limitation, we developed an AHS to grow the obligate parasite *C. campestris* without a plant host. The AHS allows growth of *C. campestris* under axenic conditions while exhibiting a parasitic habit including haustorial function. This enables studies of *Cuscuta* that more fully approximate natural mechanisms of nutrient acquisition and growth of the parasite. Additionally, the AHS supports production of viable seeds and new shoots under *in vitro* conditions (Fig. **2**), which may be useful in the genetic manipulation of *Cuscuta*.

*Cuscuta* will coil around almost any solid object of the appropriate size, shape, and orientation, which is usually anything resembling a vertical plant stem (Furuhashi *et al.*, 2011). During the optimization of the AHS, we tested different diameters and types of artificial host, including paper straws, rolled filter papers, and solid paper spindles. The best substrates were cylindrical with a relatively small diameter and a firm structure. The material also must deliver water and nutrients to the parasite. We tested liquid and solid media inside straws, but ultimately recognized that substrates with capillary capacity provided sufficient delivery of liquid media to the parasite. It is possible that other materials can be used for artificial hosts, but we found these to be simple and effective.

Haustoria structures developed in the AHS appear to function in uptake similar to natural haustoria. In this system, *C. campestris* interacts with the artificial host to take up water and small molecules as demonstrated by the uptake of CFDA and phytohormones (Fig. **3**, **4**, **6**, **7**). The CFDA uptake and translocation provided evidence of a symplastic route from the haustorium structure to the shoot tips of *Cuscuta*. Gene expression in AHS-grown haustorial regions also shows an alignment with its function in nutrient uptake and interaction with hosts, with similarities to genes induced in haustoria of parasites on actual plant hosts (Ranjan *et al.*, 2014). For example, we found that haustorial regions had enhanced expression of genes categorized as associated with increased response to stress, catabolism, and transport (carbohydrates, amino acids and ions), but reduced expression for morphogenesis and cell wall associated genes (Fig. **8**). Haustorial regions also showed activation of genes involved in response to abiotic and biotic stress.

### Insights into parasitism

The AHS provides a system in which to explore other aspects of *Cuscuta* biology. For example, what is the nature of host specificity for *C. campestris* when it grows relatively normally on a non-living host? *Cuscuta* species are recognized as having broad host ranges (Mishra, 2009), but the bases for host specificity are uncertain and have been variously attributed to host anatomy (Dawson *et al.*, 1994), host chemical composition (Honaas *et al.*, 2013), or perhaps some type of passive or active defense mechanism (Kaiser *et al.*, 2015). The AHS described herein suggests that a minimum host provides water, sugar, mineral nutrients and at least one cytokinin. It should be recognized that *Cuscuta* in the AHS is not equivalent to one growing on a preferred plant host in terms of vigor, so the AHS lacks certain aspects of the parasite-host interaction that may contribute to optimal parasite growth. But the ability of the parasite to subsist and produce viable seeds in such a system serves as a strong reminder that this parasite is extremely adaptable and can complete its life cycle with few resources.

A notable aspect of the AHS is how well the haustorial development appears to parallel that of normal haustoria on living host plants. We found evidence for an effect of the hormones NAA and BA on the parasite, with holdfast phenotype (Fig. **4**) and parasite growth (Fig. **6**) responding to hormone treatment applied through the artificial host. Elongation of cells at the holdfast surface agree with reports on the effect of injected auxin in parasite tissues (Löffler *et al.*, 1999). On the other hand, our results contrast with those of another system for artificial haustoria induction where no visible alterations in haustoria structures were observed after 48 hours in contact with a solid media containing phytohormones and without contact with the host tissues (Kaga *et al.*, 2020). We postulate that the shorter time allowed for haustorial development or different media composition could explain the conflicting results.

The addition of the cytokinin BA to the media had a pronounced effect on parasite development and growth. Parasites in the AHS with BA showed more robust holdfast morphology and greater biomass accumulation, as well as a capacity to form shoots capable of making new host connections and setting viable seeds (Fig. **4**,**6**,**7**). Gene expression data showed “response to cytokinin” as an enriched process in parasite shoots as compared to haustoria (Fig. **8b**). In a previous study, BA applied to cut ends of *C. reflexa* stem sections induced haustoria in the absence of the host and in darkness, with the maximum number of individual haustoria formed close to the apex (Ramasubramanian *et al.*, 1988). Although we did not monitor the number of individual haustoria (holdfasts in our case), we did not note differences in the capacity to form holdfasts in parasites receiving diverse treatments such as water or media, including media with BA (Fig. **4a**).

Another example of the minimum host requirements for *C. campestris* growth is the formation of vascular tissue in the haustorial structure. The development of the *C. pentagona* xylem has been correlated with differentiation of haustorial searching hyphae that contact host xylem (Vaughn, 2006). In the AHS, tracheary elements formed in the haustorial structure in contact with host-derived auxin (NAA) and cytokinin (BA), sugar (glucose) and other nutrients in the media, suggesting that these are sufficient to induce differentiation in at least some cells (Fig. **5**). This agrees with the crucial role for auxin and cytokinin in vascular differentiation (Wetmore & Rier, 1963; Aloni, 1987) and vascular reconnection during formation of grafting unions (Melnyk *et al.*, 2015). Additionally, our observation of xylem elements is supported by transcriptional data that show an enrichment in xylem development and lignin biosynthetic process in haustorial regions compared to stems 15 dpi (Fig. **8a**). Kaga et al. (2020) report the expression of genes related to xylem differentiation in *C. campestris* at 57 hours post inoculation without penetrating a plant host, indicating that haustoria acquire potential for xylem differentiation at early stages.

Other topics where the AHS may provide insight include defense and reproduction. We found genes related to defense induced in the haustorial region relative to the stem, which may indicate a preparation of the parasite for the biological interaction. It is clear that a complex molecular interaction occurs between parasite and host (Hegenauer *et al.*, 2016, 2020), leading to outcomes such as host resistance, or parasite success (Hegenauer *et al.*, 2016; Jhu *et al.*, 2019). The AHS may be useful in dissecting aspects of host and parasite defenses. The issue of flowering is another exciting topic, with recent work suggesting that the parasite relies on its host for regulation of flower induction (Shen *et al.*, 2020). Some have contended that *C. campestris* has lost many of its own flowering regulator genes through evolutionary reduction (Vogel *et al.*, 2018), so it must therefore rely on flowering signals from a host plant that serve to synchronize its flowering times with the host. However, we observed parasites flowering and producing seeds in the absence of a flowering – or even a living - plant host. Further study is needed to understand flowering dynamics in *Cuscuta*, and the AHS may be a useful tool.

Historically microbial cultures and especially axenic cultures have allowed a deeper understanding of microorganisms and parasites biology. For example, the growing of human parasites *in vitro* has enabled the study of their biological features and requirements without the influence or contamination of the host. It has also facilitated their identification and the assessment of control methods (Visvesvara & Garcia, 2002). The AHS presents a valuable new approach to studying *Cuscuta*. In this system, it is possible to manipulate “host” chemical composition such as phytohormones (e.g., Fig. **4**,**6**,**7**), or to test hypothesis involving environmental conditions that would otherwise be impractical with a live host (e.g., growth in darkness). The AHS is versatile and may be modified to explore different substrate material or compare specific host requirements of different *Cuscuta* species, although the AHS was optimized for *C. campestris*. We expect this tool to be a valuable addition to the workbench of *Cuscuta* researchers.

## Acknowledgements

Hannah Ambrose helped with material preparation for AHS experiments.

## Author contributions

VBG and JHW conceived the project. VBG developed the system, conducted the experiments, analyzed the data and drafted the manuscript. JHW edited the manuscript.

## Data availability

The data that support the findings of this study are openly available in the NCBI Gene Expression Omnibus (GEO) archive at https://www.ncbi.nlm.nih.gov/geo/query/acc.cgi?acc=GSE178396, reference number GSE178396.

## Declarations

- Ethics approval and consent to participate Not Applicable
- Consent for publication Not Applicable
- Competing interests The authors declare that they have no competing interests.

## Supporting Information

**Fig. S1** Materials for building the artificial host system (AHS).

**Fig. S2** *C. campestris* with developed turgid holdfast gained fresh weight, biomass and length compared to parasites with no developed turgid holdfast.

**Fig. S3** New parasite shoots produce in the AHS are viable.

**Fig. S4** Relationship among biomass, length, and fresh weight in *C. campestris*.

**Fig. S5** Relationship among biomass, length, and flowers and fruits presence in *C. campestris.*

**Fig. S6** Differential gene expression analysis on haustorial region and stem tissues collected from *C. campestris* growing in the AHS.

**Fig. S7** Comparison between up regulated genes present in three categories of haustoria developmental conditions.

**Fig. S8** Singular enrichment analysis (SEA) of *C. campestris* up regulated genes in haustorial regions under different conditions.

**Table S1** Enriched GO terms for upregulated and down regulated genes.

**Table S2** DEGs shared by datasets considering haustoria functional status.

