## Supplementary Figures for "An artificial host system enables the obligate parasitic plant *Cuscuta campestris* to grow and complete its life cycle *in vitro*"

The following Supporting Information is available for this article:

**Fig. S1** Materials for building the artificial host system (AHS).

**Fig. S2** *Cuscuta campestris* with developed turgid holdfast gained fresh weight, biomass and length compared to parasites with no developed turgid holdfast.

**Fig. S3** New parasite shoots produce in the artificial host system (AHS) are viable.

**Fig. S4** Relationship among biomass, length, and fresh weight in *Cuscuta campestris*.

**Fig. S5** Relationship among biomass, length, and flowers and fruits presence in *Cuscuta campestris*.

**Fig. S6** Differential gene expression analysis on haustorial region and stem tissues collected from *Cuscuta campestris* growing in the artificial host system (AHS).

**Fig. S7** Comparison between up regulated genes present in three categories of haustoria developmental conditions.

**Fig. S8** Singular enrichment analysis (SEA) of *Cuscuta campestris* up regulated genes in haustorial regions under different conditions.

**Table S1** Enriched Gene ontology (GO) terms for upregulated and down regulated genes

**Table S2** Differentially expressed genes (DEGs) shared by datasets considering haustoria functional status

**Fig. S1** Materials for building the artificial host system (AHS). Cotton is removed from one end of a swab to yield a 0.3 cm diameter, 7 cm long paper spindle, the spindle is held vertical by a thin sheet of plastic, with the cotton end directed towards the bottom of the box to be immersed in media. Four "artificial host" spindles are thus held together and placed in a culture box.

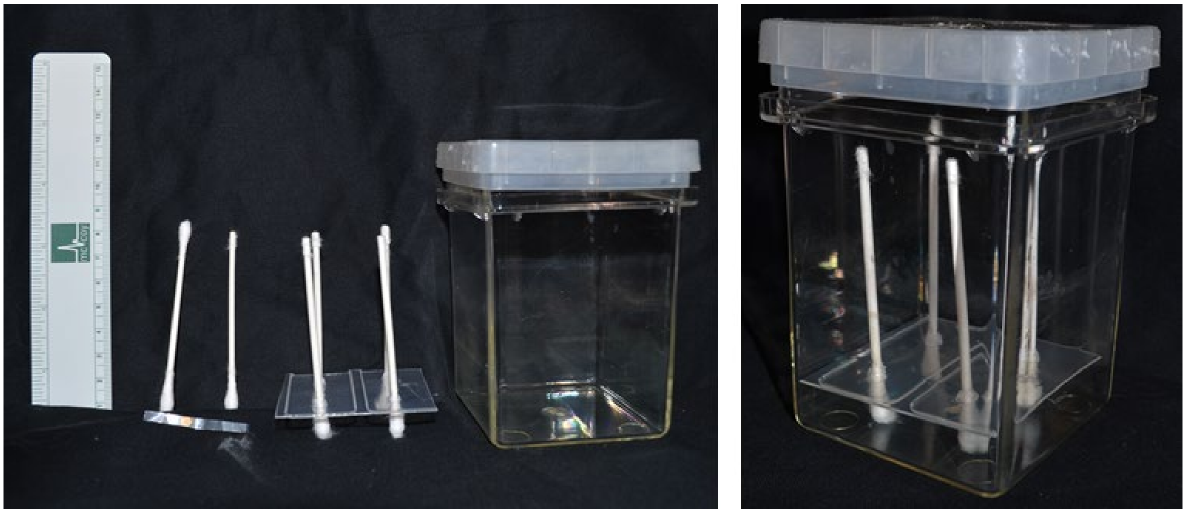

**Fig. S2** *C. campestris* with developed turgid holdfast (Hf) growing in the artificial host system (AHS) with MMS (Modified Murashige and Skoog) media including 1-naphthaleneacetic acid (NAA) (3 mg $l^{-1}$) and 6-benzylaminopurine (BA) (1 mg $l^{-1}$) gained fresh weight (**a**), biomass (**b**) and length (**c**) compared to parasites with no developed turgid holdfast (No Hf) under the same media, or under water as media or without any media ("nothing"). Parasites growing without any media did not present developed turgid holdfast or growth. Evaluation was made at 36 days post inoculation (dpi). Statistical differences were detected by ANOVA with post-hoc Tukey HSD test. Differences were considered statistically significant at P<0.05 and indicated with different letters. Number of samples ranged from 10 to 51 per treatment. Data correspond to three independent experiments.

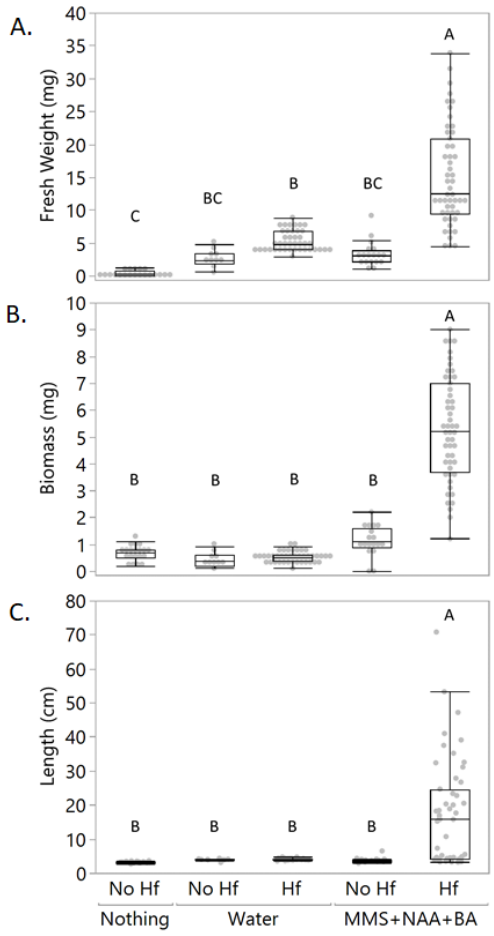

**Fig. S3** New parasite shoots produce in the artificial host system (AHS) are viable. **a.** *Cuscuta campestris* new shoots obtained through the AHS using a 3cm shoot tip from nursery as initial inoculum 36 days post inoculation. Orange arrow shows an active shoot. Blue arrow shows a senescent shoot. **b**. Active shoots were able to colonize a new plant host (*Arabidopsis thaliana*).

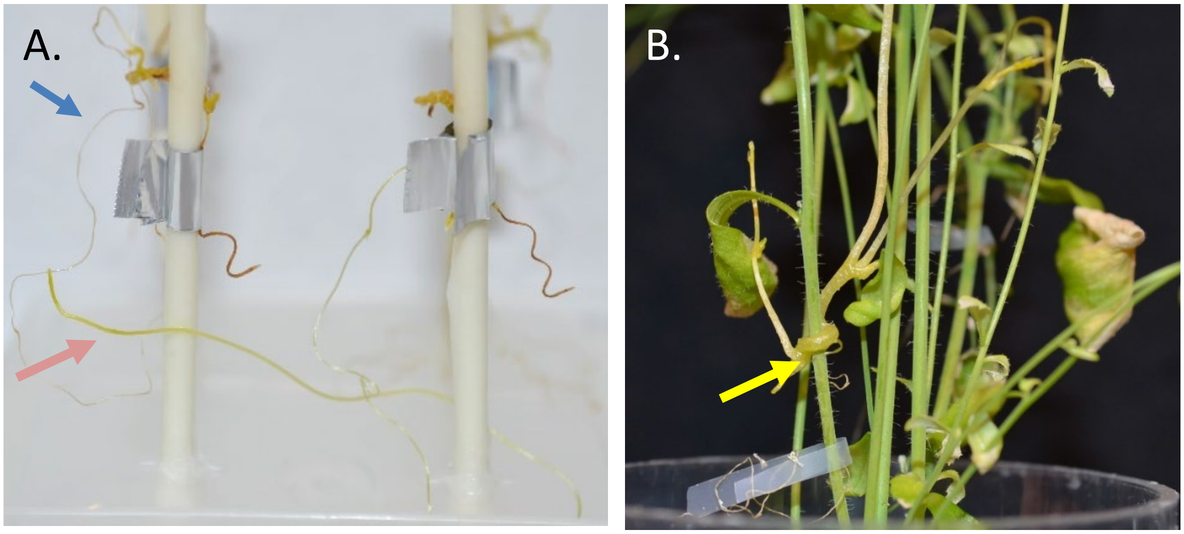

**Fig. S4** Relationship among biomass, length, and fresh weight in *Cuscuta campestris*. Graphic collects data from parasites growing under different treatments in the artificial host system (AHS) 36 dpi. Pearson's coefficient (r) was calculated regardless of treatments. Colors correspond to different treatments and are shown only for reference. Ellipses indicate the area where 95% of the data fall. n= 210. Stars indicate the mean point of the ellipse and correspond total average. Data correspond only to parasites that coiled and developed holdfast.

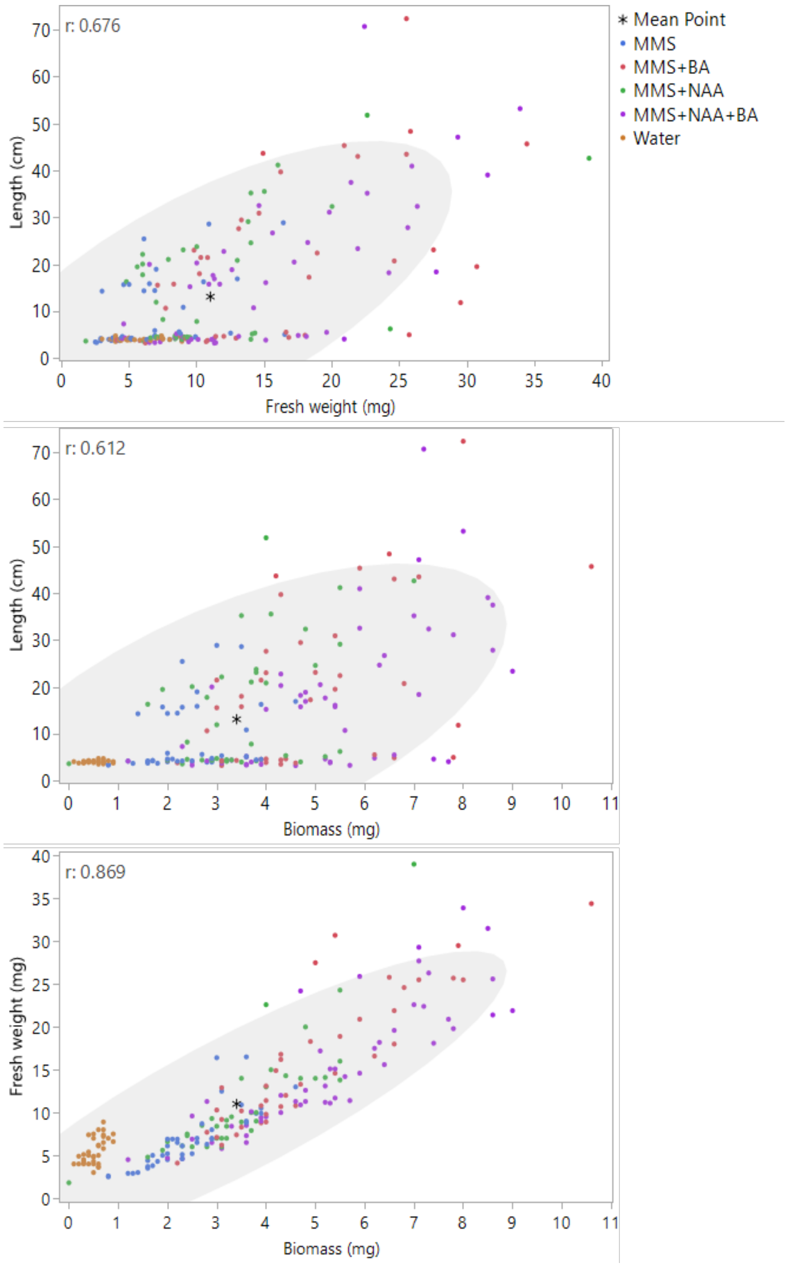

**Fig. S5** Relationship among biomass, length, and flowers and fruits presence in *Cuscuta campestris.* Graphic collects data from parasites growing under different treatments in the artificial host system (AHS) 36 days post inoculation (dpi). Ellipses indicate the area where 50% of the data corresponding to a specific condition fall: parasites with no flowers or fruits (grey, n=147), parasites with only flowers (orange, n=50), parasites with flowers and fruits (blue, n=13). Stars represent the average for the values in each ellipse. Data correspond only to parasites that coiled and developed a turgid holdfast.

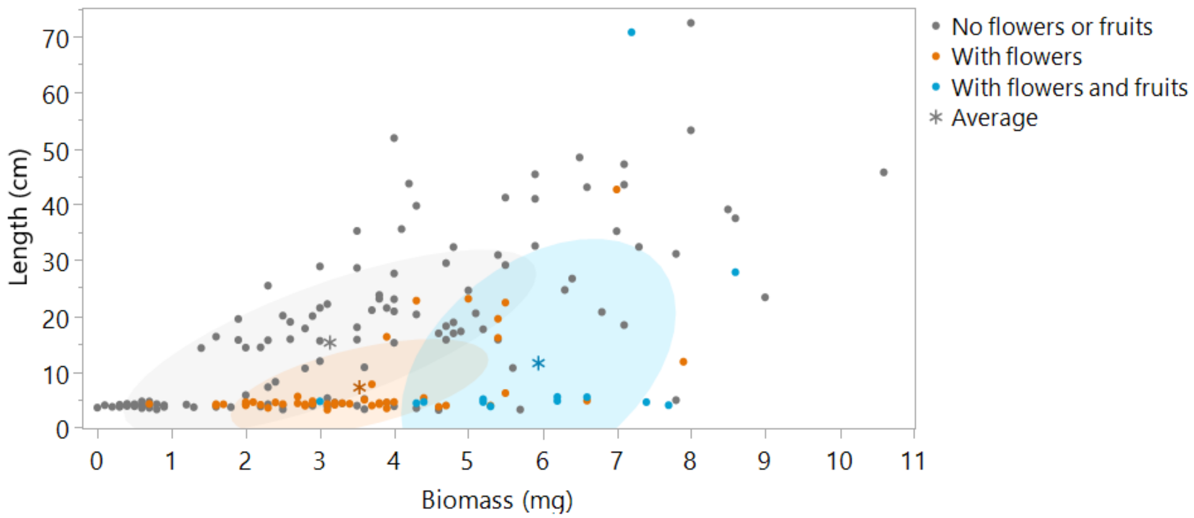

**Fig. S6** Differential gene expression analysis on haustorial region and stem tissues collected from *Cuscuta campestris* growing in the artificial host system (AHS). Principal component analysis (PCA) shows a segregation on the samples by type of tissue. Differential expression analysis results show the number of down regulated or up regulated genes in the haustorial region tissues compare to the stem tissues after filters were applied. Four biological replicates of each tissue were sequenced. Illumina RNA-seq yielded an average of 63.4 million total raw reads per sample and, after filtering by length and low quality, 97.8% passed quality control. False discovery rate: FDR. Adjusted p-value: padj.

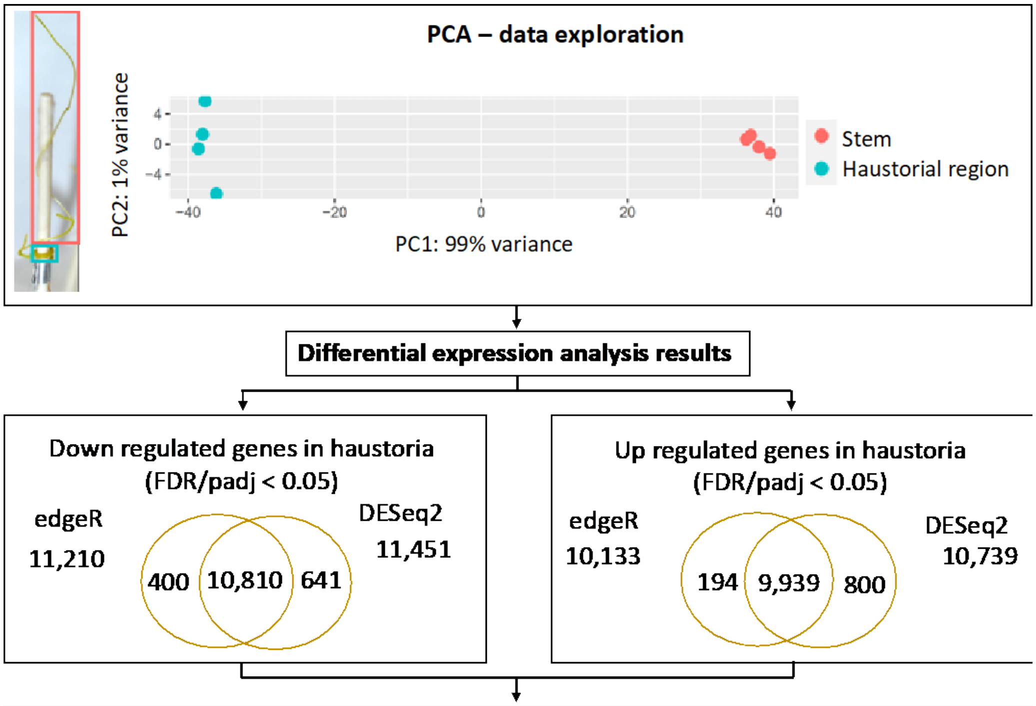


| Filter | Total DEG | Up regulated in Haustoria | Down regulated in Haustoria |
|---|---|---|---|
| padj < 0.05 | 20,749 | 9,939 | 10,810 |
| padj < 0.01 | 11,479 | 8,969 | 2,510 |

**Fig. S7** Comparison between up regulated genes present in three categories of haustoria developmental conditions. First category corresponds to up regulated genes (8.969 genes, padj<0.01) in the functional haustorial regions of parasites growing in artificial host system (AHS). The second category includes independently reported genes expressed in functional haustorial region of *C. campestris* growing on different plant hosts. Out of the 469 genes in this group, 177 correspond to genes reported by Kim et al (2014) and 308 to Ranjan et al (2014), of which 16 were shared by both studies. The third category corresponds to up regulated genes in haustorial tissue, 87 hours after the artificial-induction of haustoria in contact with a leave of a plant host (Kaga *et al.*, 2020). 220 genes were shared between all categories, with haustoria formation regardless function or contact with plant host. 62 genes were shared between the groups that harbored functional haustorial regions regardless the presence of a plant host. While 78 genes were shared only between the groups that had a contact with plant host tissues regardless haustorial function. Adjusted p-value: padj.

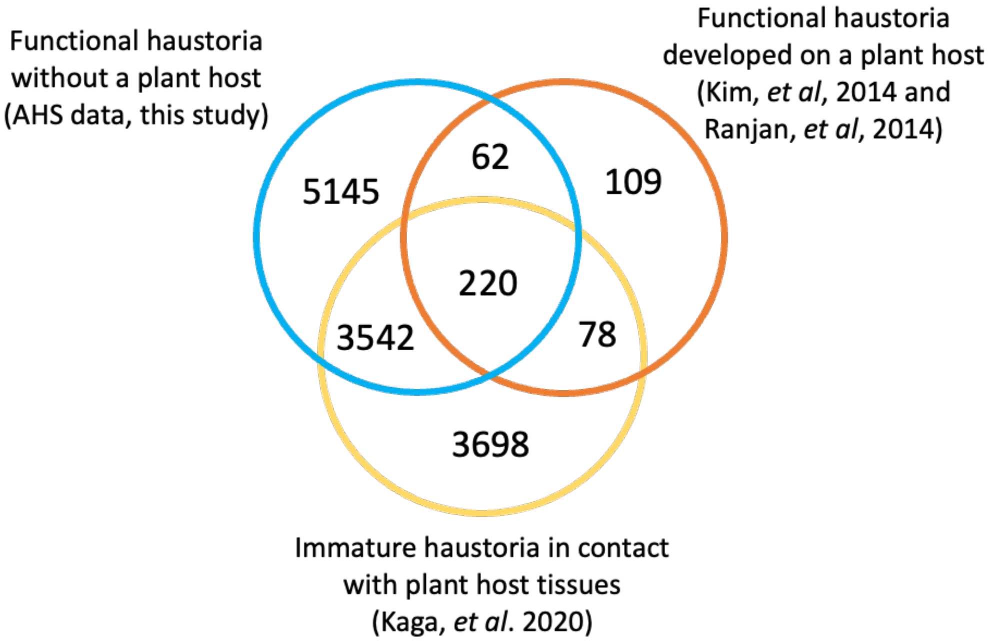

**Fig. S8** Singular enrichment analysis (SEA) of *Cuscuta campestris* up regulated genes in haustorial regions under different conditions. Gene ontology (GO) Slim terms are based on homologue candidate genes in *Arabidopsis thaliana* and SEA was made considering *A. thaliana* genome as reference. GO Slim plant terms are organized in a hierarchical way and show **a**. GO terms associated to genes shared between datasets that include functional and immature haustoria regardless contact with a host plant, and **b**. GO terms associated to genes shared between datasets that include only functional haustoria regardless contact with a host plant. A functional haustoria is determined by formation of conspicuous holdfasts, and growth and development of new shoots and flowers after haustoria formation.

A.

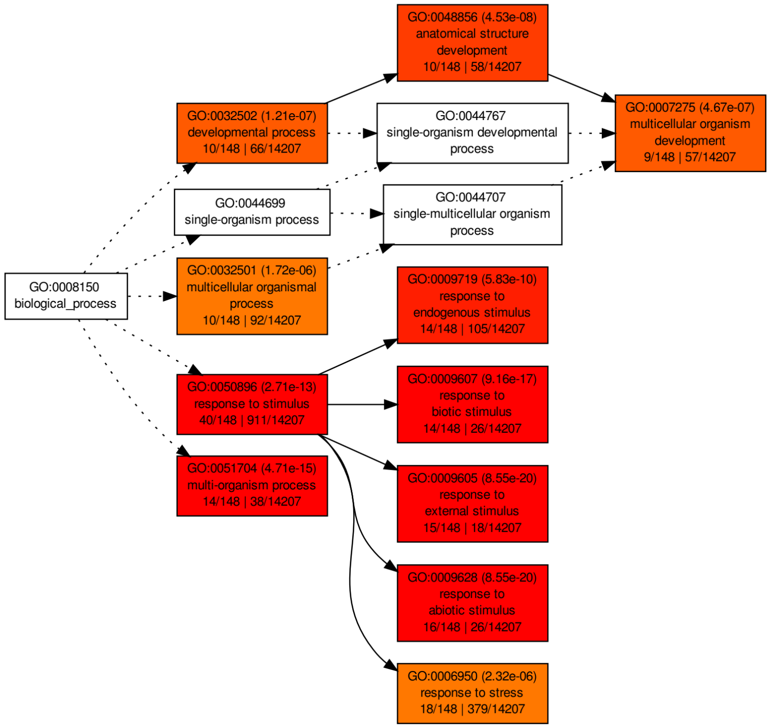


B.

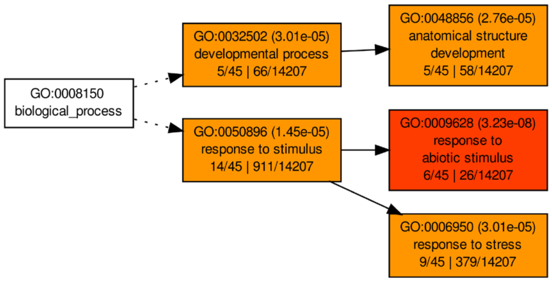
